# Homoharringtonine targets XPC to inhibit DNA damage repair and sensitize tumors to platinum-based chemotherapy

**DOI:** 10.64898/2026.10.27.762176

**Authors:** Chuanjie Wang, Ying Zhang, Xiaoqin Zhou, Jia He, Longjiang Liu, Luyi Wang, Yanfang Zhang, Qin Gong, Jiayu Zhang, Yongjun Dang, Jiaxue Wu

**Author notes:** Corresponding author. E-mail addresses (Jiaxue Wu); (Yongjun Dang). These authors contributed equally (Chuanjie Wang, Ying Zhang, Xiaoqin Zhou).

## Abstract

DNA damage repair (DDR) is essential for preserving genomic stability and also represents a key targetable axis in cancer drug development. Here, we identify and validate homoharringtonine (HHT), a clinically approved antileukemic drug, as a novel DDR inhibitor using a phenotype-driven, high-content screen based on 53BP1 Tudor-domain foci formation. Mechanistically, HHT directly binds XPC, a critical nucleotide excision repair (NER) factor, with high affinity (Kd = 158.4 nM) in a manner dependent on Thr795 of XPC. HHT disrupts XPC–damaged DNA binding, impairs NER, and reduces non-homologous end joining (NHEJ) efficiency. Consequently, HHT sensitizes tumor cells to platinum-based chemotherapy in vitro and in vivo. These findings establish a phenotype-based platform for DDR inhibitor discovery, identify HHT as a first-in-class XPC inhibitor and uncover a previously unrecognized mechanism of HHT that may be exploited to enhance platinum-based therapy in solid tumors.

## Introduction

The DNA damage repair (DDR) machinery serves as the primary guardian against constant insults from both endogenous metabolism and exogenous genotoxic agents^1^. DDR also plays a critical and dual-faceted role in tumor progression and the development of resistance to radiotherapy and chemotherapy^2–4^. Consequently, DDR has emerged as a compelling and targetable axis in oncology^5^.

Despite substantial progress in DDR-targeted drug discovery over the past decades and a variety of small-molecule DDR inhibitors identified through conventional target-based screening have advanced into early-phase clinical trials, the clinical translation of such agents has proven challenging^6–8^. To date, inhibitors of poly(ADP-ribose) polymerase (PARP) remain the only class of DDR-targeted therapeutics to have achieved regulatory approval and established a firm foothold in routine clinical practice, primarily in the context of homologous recombination-deficient tumors^9^. Other inhibitors targeting ATM^10^, ATR^11^, CHK1/2^10^, DNA-PK^12^, WRN^13,14^, USP1^15^, POLQ^16^, and WEE1^17^ remain under active investigation, with numerous preclinical and early-phase clinical studies exploring their potential. This disparity underscores the pressing need to discover novel DDR inhibitors with distinct mechanisms of action, broader application spectra, or the capacity to circumvent acquired resistance, as well as to explore alternative screening strategies beyond conventional target-centric approaches. Among the various DDR subpathways, nucleotide excision repair (NER) plays a critical role in removing bulky DNA adducts, including those induced by platinum-based chemotherapeutics^18^. The efficiency of NER directly influences tumor cell sensitivity to platinum-based drugs, and elevated expression and enhanced activity of NER-related proteins are well-recognized as major contributors to platinum resistance^4^. Therefore, targeted inhibition of the NER pathway represents a core strategy to potentiate platinum-based chemotherapy and overcome drug resistance. Despite the significant clinical value of targeting NER, the development of NER inhibitors has lagged considerably behind that of inhibitors targeting other DDR pathways^19^. The primary reason is that the precise operation of the NER pathway heavily relies on protein-protein interactions (PPIs)—such as the formation of the ERCC1/XPF heterodimer and the cooperative binding of XPA with other NER factors—rather than on key enzymes with well-defined catalytic activities, as is the case for most DDR pathways^20^. Compared with the traditional "lock-and-key model" (i.e., enzyme and its inhibitor), using small molecules to disrupt PPIs is more challenging in medicinal chemistry, as these interactions involve large and relatively flat binding interfaces^21^. To date, no drug targeting the NER pathway has been approved globally, and only a limited number of small-molecule compounds, PROTAC degraders, or molecular glues targeting the interactions of XPA^22^, XPB, XPG^23^, and ERCC1/XPF^24^ are in preclinical research stages.

Here, we performed a phenotype-driven, high-content screen based on 53BP1 Tudor-domain foci formation and identified HHT as a novel DDR inhibitor. We demonstrated that XPC, a critical sensor and initiator of the NER pathway, serves as the direct and physiologically relevant molecular target of HHT in solid tumors. HHT binds to the DNA-binding region of XPC, thereby disrupting the interaction between XPC and damaged DNA and sensitizing tumor cells to platinum-based chemotherapy. These results identify HHT as a first-in-class XPC inhibitor that can be used as a platinum-sensitizing agent in the treatment of solid tumors, thereby broadening the therapeutic scope of this clinically established drug and offering a promising strategy to overcome platinum resistance in oncology.

## Results

### High-throughput phenotypic screening identifies homoharringtonine as a potent inhibitor of DNA damage repair

To identify small molecules capable of suppressing DNA damage repair, we established a high-content phenotypic screening platform based on 53BP1 Tudor-domain foci formation. 53BP1 is a downstream factor in the DNA repair pathway and serves as a well-established marker for monitoring repair activity. A U2OS cell line stably expressing EGFP-Tudor was generated, in which bleomycin treatment induced robust and readily quantifiable nuclear foci formation (Fig. S1A). After confirming a sufficiently broad dynamic range between untreated and bleomycin-treated cells, we optimized both the concentration and duration of bleomycin exposure to maximize assay performance. The ATM inhibitor KU-55933 was used as a positive control, and evaluation of the assay robustness demonstrated an acceptable Z′-factor (0.542) for high-throughput screening (Fig. S1B).The overall screening workflow is illustrated in Fig.S1C. Briefly, EGFP-Tudor-U2OS cells were seeded 24 hour prior to compound treatment. Candidate compounds were added and incubated for 1 hour, followed by bleomycin treatment for an additional 2 hour to induce DNA damage. Cells were subsequently subjected to automated image acquisition using a high-content imaging system, and Tudor-foci formation was quantified as the primary screening readout.

Using this platform, we screened a natural product library of 2,592 small molecules (L6000, TargetMol) and identified 25 compounds that significantly inhibited bleomycin-induced Tudor-foci formation in the primary screen (Fig. 1A). Secondary screening of these 25 candidates under concentration-gradient conditions revealed that 11 compounds retained significant inhibitory activity at concentrations as low as 1 μM (Fig.1B). These 11 compounds were subjected to a third round of concentration-response validation, which identified 9 compounds that inhibited Tudor-domain foci formation in a concentration-dependent manner.(Fig.1C). Among the 9 identified hits, homoharringtonine (HHT) and its structural analog harringtonine (HT) emerged as the most attractive candidates(Fig.1D). HHT is a natural alkaloid originally developed in China and widely used for hematological malignancies, especially chronic myeloid leukemia and acute myeloid leukemia^25,26^. While its anti-leukemic activity has been extensively characterized, its direct targets and mechanisms in solid tumors, especially regarding DNA damage repair, remain largely unknown. Our screening results uncover a previously unrecognized role for HHT in DDR regulation and provide a foundation for exploring its underlying molecular mechanism.

**Fig. 1.**
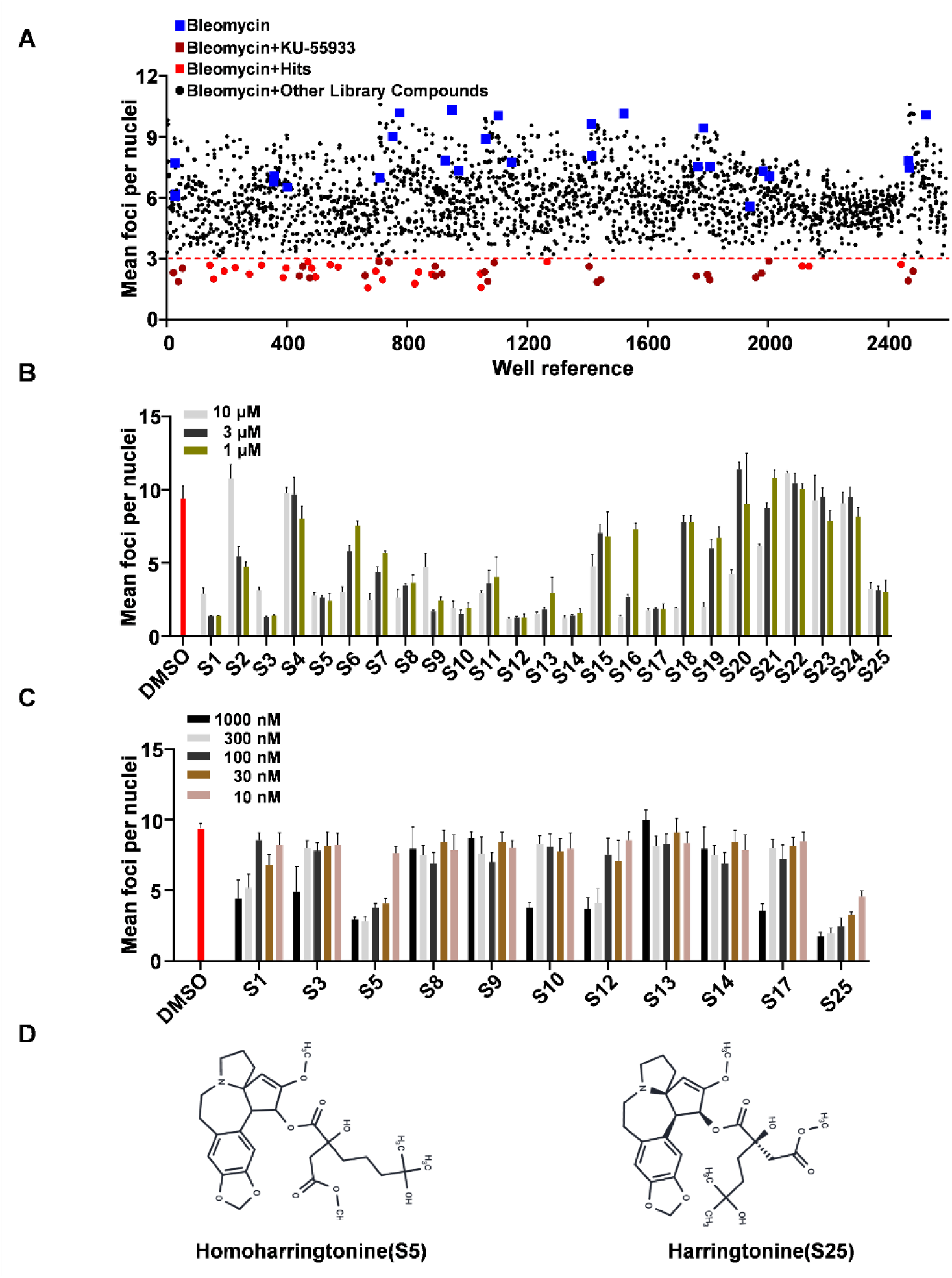
Identification of compounds that inhibit DNA damage repair by high-content screening (HCS). (A) Primary screening of a natural product library containing 2,592 small molecules identified 25 candidate compounds that effectively suppressed EGFP-Tudor foci formation at a concentration of 10 μM. (B) The 25 primary hits were subjected to a first round of dose-dependent rescreening at concentrations ranging from 10 μM to 1 μM. 11 compounds with stronger inhibitory activity were selected for further validation. (C) The 11 selected candidates were further evaluated in a second round of dose-dependent rescreening at concentrations ranging from 1,000 nM to 10 nM,S5 and S25 indicate Homoharringtonine and harringtonine respectively. (D) Chemical structures of homoharringtonine and harringtonine.

### HHT compromises DNA damage repair

Given its well-established clinical safety profile and FDA approval, HHT was selected for subsequent mechanistic investigation. We first assayed the half-maximal inhibitory concentration (IC₅₀) of HHT for suppressing Tudor-domain foci formation and determined it to be 18.8 nM (Fig. 2A). Interestingly, although HHT suppressed bleomycin-induced Tudor-domain and endogenous 53BP1 foci formation, it had no effect on the DNA damage marker γH2AX foci formation(Fig. 2B-C), suggesting that HHT acts downstream of H2AX in the DNA damage response pathway. Although HHT is canonically known to inhibit ribosomal function and thereby attenuate global protein synthesis, we observed that the expression levels of 53BP1 and EGFP-Tudor remained unchanged upon HHT treatment in our experimental conditions (Fig. S2), indicating that its effect on foci formation is not due to reduced protein abundance. Furthermore, we found that HHT treatment delayed the recovery of γH2AX level induced by bleomycin in multiple tumor cell lines including U2OS, CT26 and HCT116(Fig. 2D and Fig. S3 ), further indicating that HHT suppresses DNA damage repair.

**Fig. 2.**
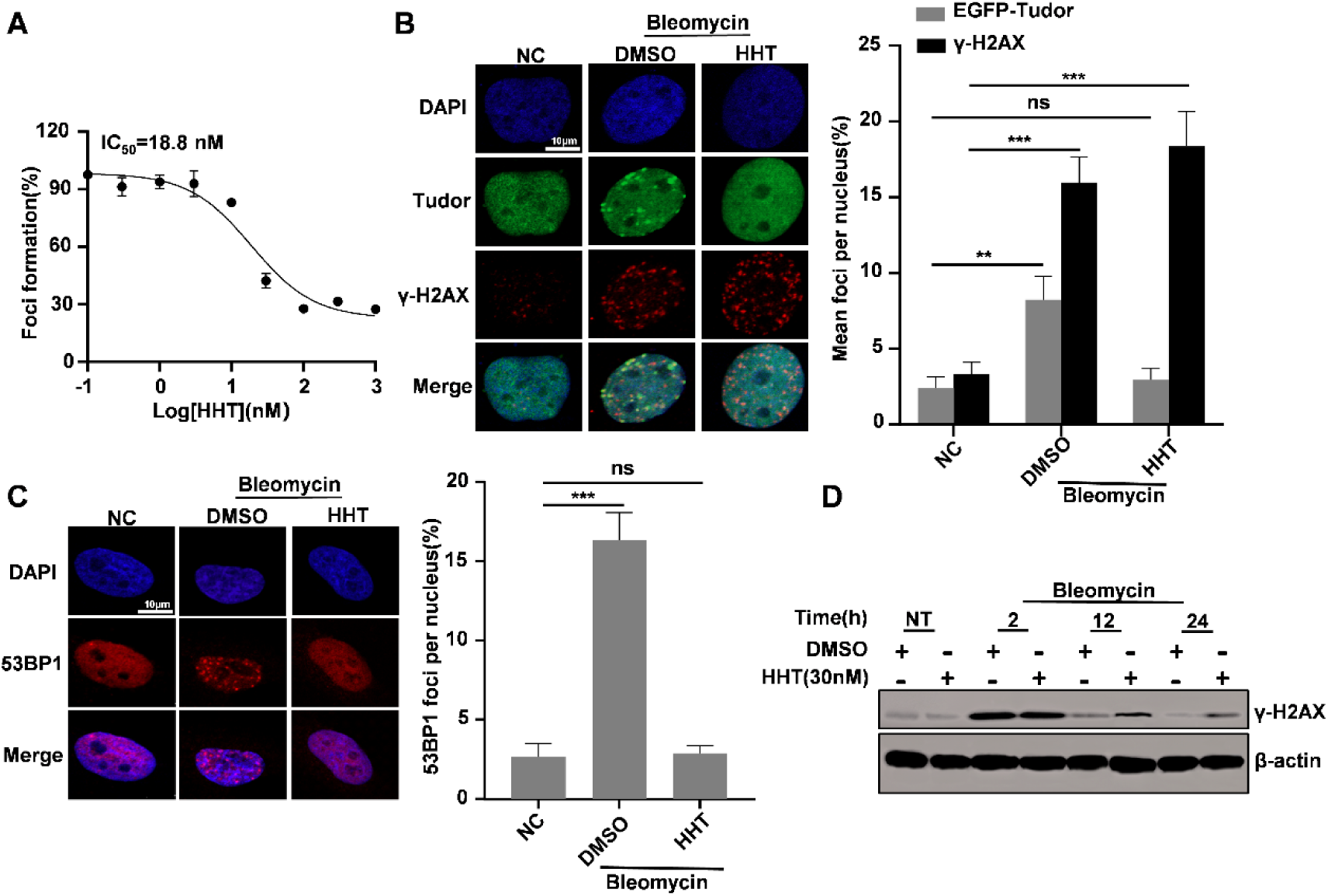
Homoharringtonine inhibits the DNA damage repair phenotype. (A) Dose-response analysis showing that HHT inhibited EGFP-Tudor foci formation, with an IC₅₀ of 18.8 nM. (B) EGFP-Tudor-U2OS cells were pretreated with HHT (30 nM) for 1 h, followed by bleomycin treatment for 2 h to induce DNA damage. Representative immunofluorescence images showing EGFP-Tudor and γ-H2AX foci formation are presented . Quantification of the mean number of EGFP-Tudor and γ-H2AX foci per nucleus is shown. (C) U2OS cells were pretreated with HHT (30 nM) for 1 h and then exposed to bleomycin for 2 h. Endogenous 53BP1 foci formation was detected by immunofluorescence, and representative images are shown. Quantification of the mean number of 53BP1 foci per nucleus is shown . (D) U2OS cells were pretreated with HHT (30 nM) for 1 h, followed by bleomycin treatment for 2 h. After bleomycin removal, cells were maintained in medium containing HHT and collected at the indicated time points. γ-H2AX levels were analyzed by western blotting.Data are presented as mean ± SEM from three independent experiments. Statistical significance was determined by one-way ANOVA for panels (B) and (C). ns, not significant; \*\**P* < 0.01; \*\*\**P* < 0.001.

### XPC is the target of HHT in DNA damage repair

To elucidate the molecular target of HHT, we first prepared HHT-conjugated beads (HHT-Beads) via photochemical crosslinking and incubated them with total protein from cell lysates to enrich interacting proteins for mass spectrometry analysis(Fig. S4) . MS profiling revealed multiple proteins specifically enriched by HHT-Beads compared with control beads, including NκRF and EWSR1, which have been previously reported as direct HHT targets^27,28^. Xeroderma Pigmentosum Complementation Group C (XPC), a core nucleotide excision repair (NER) factor, was specifically enriched in the HHT-Beads complex(Fig.3A). Notably, XPC was also present in previous HHT affinity purification datasets^27^, although it was not investigated further in that study. Furthermore, previous studies indicate that XPC can modulate UV-induced RNF8 foci formation^29^, which is upsteam of 53BP1, although the precise mechanism remains unclear. Based on these chemoproteomic findings and literature evidence, we hypothesized that XPC is the direct target of HHT and mediates its inhibitory effect on DNA damage repair.

**Fig. 3.**
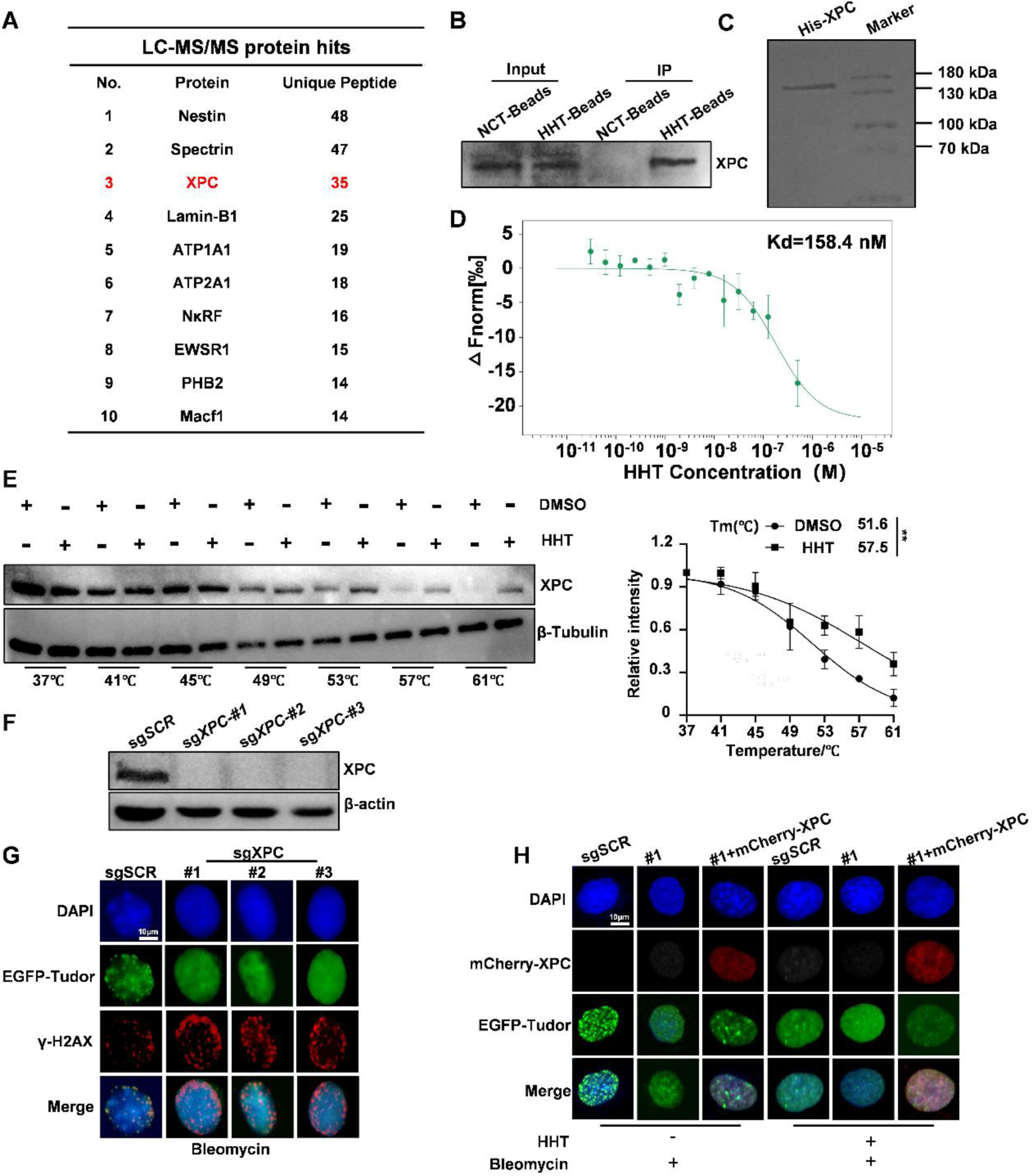
HHT directly binds XPC to inhibit DNA damage repair. (A) LC-MS/MS analysis of proteins enriched by HHT-Beads. The top-ranked candidate proteins and their corresponding unique peptide numbers are shown. XPC was identified as one of the specifically enriched proteins. (B) HHT-Beads were incubated with U2OS cell lysates, and the enrichment of endogenous XPC was detected by immunoblotting. (C) Sf9 expression system for expressing and purifying XPC protein - Coomassie brilliant blue staining. (D) Microscale thermophoresis (MST) analysis showing the binding affinity between purified XPC protein and HHT. The dissociation constant (Kd) was calculated as 158.4 nM. (E) U2OS cells were treated with HHT (80 μM) for 5 h and then subjected to a temperature-gradient cellular thermal shift assay. XPC thermal stability was analyzed by western blotting , and melting temperature (Tm) values were calculated from three independent experiments. (F) Western blot analysis confirming CRISPR/Cas9-mediated knockout of XPC in U2OS cells. (G) EGFP-Tudor-U2OS cells with or without XPC knockout were treated with bleomycin to induce DNA damage. Representative immunofluorescence images showing EGFP-Tudor and γ-H2AX foci formation are presented. (H) Reintroduction of mCherry-XPC in EGFP-Tudor-U2OS cells without XPC were pretreated with or without HHT followed by bleomycin to induce DNA damage. Representative immunofluorescence images showing EGFP-Tudor foci formation are presented.

To validate whether HHT directly binds XPC, HHT-Beads or control beads were incubated with U2OS cell lysates, followed by Western blot analysis using an anti-XPC antibody. As shown in Fig.3B, XPC was specifically enriched by HHT-Beads, not control beads. We then expressed and purified recombinant XPC protein using a baculovirus system in Sf9 insect cells (Fig.3C)and measured its binding affinity to HHT by microscale thermophoresis (MST). As shown in Fig. 3D, HHT bound to XPC with high affinity (Kd = 158.4 nM). Furthermore, a cellular thermal shift assay (CETSA) in U2OS cells treated with HHT across a temperature gradient revealed increased thermal stability of XPC, with the melting temperature (Tm) shifting from 51.6°C to 57.5°C (Fig. 3E). Next, we generated an XPC-knockout U2OS cell line (*XPC⁻/⁻*-U2OS) using CRISPR/Cas9 to examine whether XPC regulate EGFP-Tudor foci formation(Fig. 3F).As shown in Fig.3G, bleomycin-induced Tudor-foci formation was abolished in XPC deficient cells, whereas γ-H2AX foci formation remained unaffected. Reintroduction of mCherry-XPC restored EGFP-Tudor foci formation upon bleomycin exposure, and this restored foci formation was again inhibited by HHT treatment(Fig.3H). Collectively, these results demonstrate that HHT binds XPC and XPC is the direct target of HHT in DNA damage repair.

### HHT dirsputed the interation between XPC and damaged-DNA through binding with the Thr795 of XPC

To predict the HHT binding pocket and identify key interacting residues, we performed molecular docking using Autodock Vina with the crystal structure of XPC (PDB: 8EBV). The docking analysis suggested that hydrogen bonds with Thr117, Ser122, and Thr795 of XPC are critical for HHT binding (Fig. 4A). To validate these findings, we expressed and purified recombinant proteins carrying a triple mutation (XPC-TST3A) or individual point mutations (XPC-T117A, XPC-S122A, and XPC-T795A) and used them in pull-down assays with HHT-Beads. As shown in Fig. 4B, XPC-T117A and XPC-S122A, but not XPC-TST3A or XPC-T795A, were specifically enriched by HHT-Beads, suggesting that Thr795 of XPC is critical for HHT binding. Consistently with that, recombinant XPC-T795A protein lost binding affinity for HHT in MST assays and did not exhibit increased thermal stability upon HHT treatment (Fig. S5A-B). Collectively, these results demonstrate that Thr795 of XPC is critical for HHT binding.

**Fig. 4.**
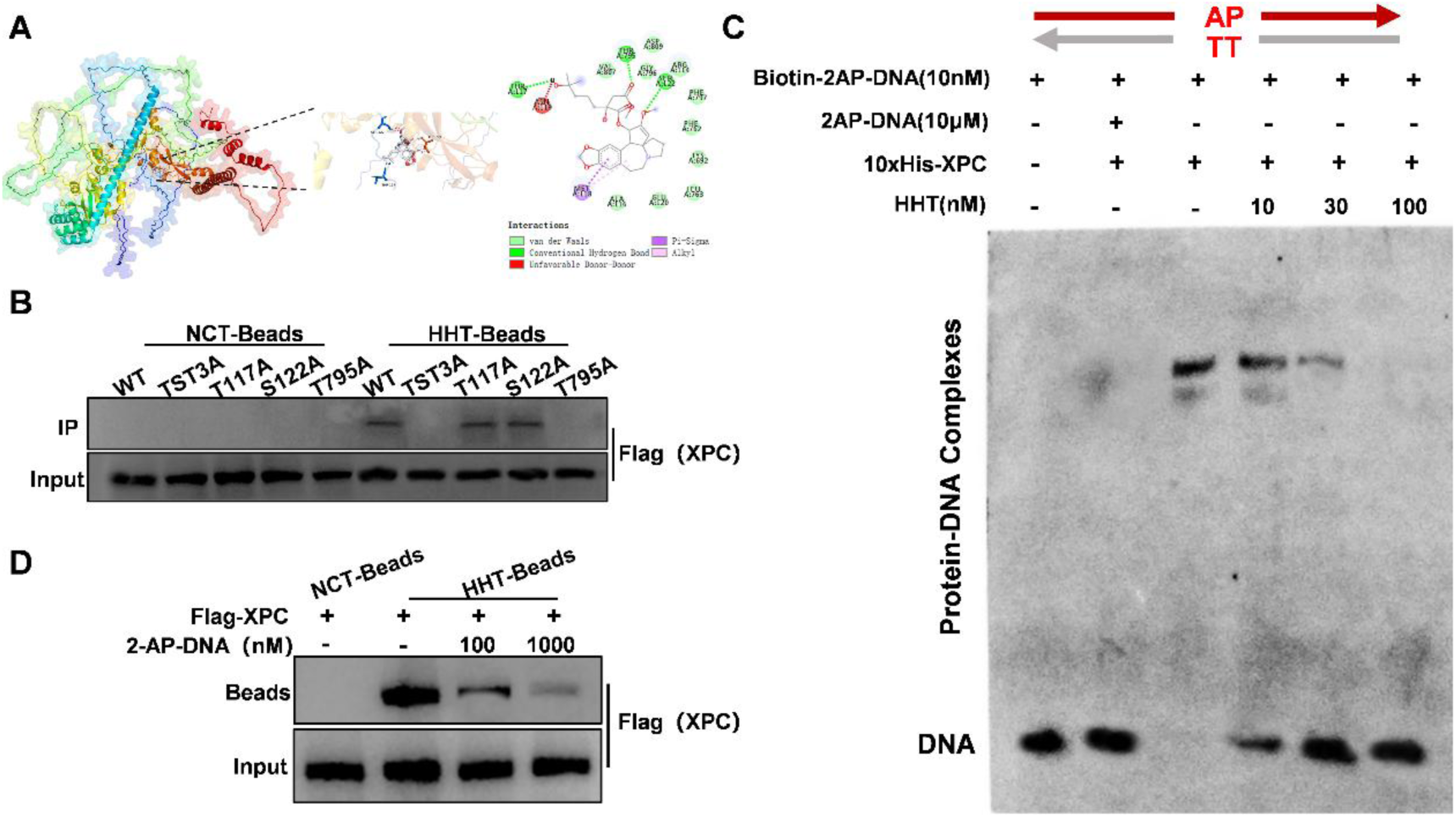
HHT binding to Thr795 disrupts XPC–damaged DNA interaction. (A) Molecular docking model showing the predicted interaction between HHT and XPC, including the putative HHT-interacting residues. (B) HEK-293T cells were transfected with plasmids encoding wild-type or mutant Flag-XPC proteins. Cell lysates were incubated with HHT-Beads, and the binding of HHT to XPC or XPC mutants was analyzed by western blotting. TST3A indicates the combined mutation of T117, S122, and T795 to alanine; T117A, S122A, and T795A indicate the corresponding single-point mutations. (C) Electrophoretic mobility shift assay (EMSA) was performed to evaluate the binding of XPC to damaged DNA in the presence of HHT. Unlabeled 2AP-DNA was used as a competitor at a 1000-fold molar excess relative to biotin-labeled 2AP-DNA. (D) XPC protein was pre-incubated with 2-AP-DNA at the indicated concentrations and then subjected to pull-down with HHT-beads. NCT-beads served as a negative control. Bead-bound XPC was detected by anti-Flag immunoblotting, with input shown as a loading control. Damaged DNA reduced XPC binding to HHT-beads in a concentration-dependent manner.

As a key lesion-recognition factor in the nucleotide excision repair (NER) pathway, XPC binds to damaged DNA, an event essential for activation of the NER machinery. Consistent with this role, XPC recognizes and binds base-deficient damaged DNA duplexes under in vitro biochemical conditions^30^. To determine whether HHT affects this DNA-binding function of XPC, we performed electrophoretic mobility shift assay (EMSA) using purified XPC protein incubated with increasing concentrations of HHT. As shown in Fig. 4C, HHT abrogated the XPC–damaged DNA interaction in a concentration-dependent manner. Consistently, pull-down assays showed that damaged DNA reduced the amount of XPC captured by HHT-Beads, indicating that damaged DNA competes with HHT-Beads for XPC binding. (Fig. 4D). Collectively, these results indicate that HHT directly compromises the DNA-binding activity of XPC.

### HHT inhibits both NER and NHEJ

Given the critical role of XPC in NER, we next examined the effect of HHT on NER activity. XPC-deficient cells exhibit markedly reduced repair capacity toward UV-C-irradiated plasmids^31^. Using this well-established system, we demonstrated that HHT inhibited cellular NER activity in a concentration-dependent manner (Fig. 5A).

**Fig. 5.**
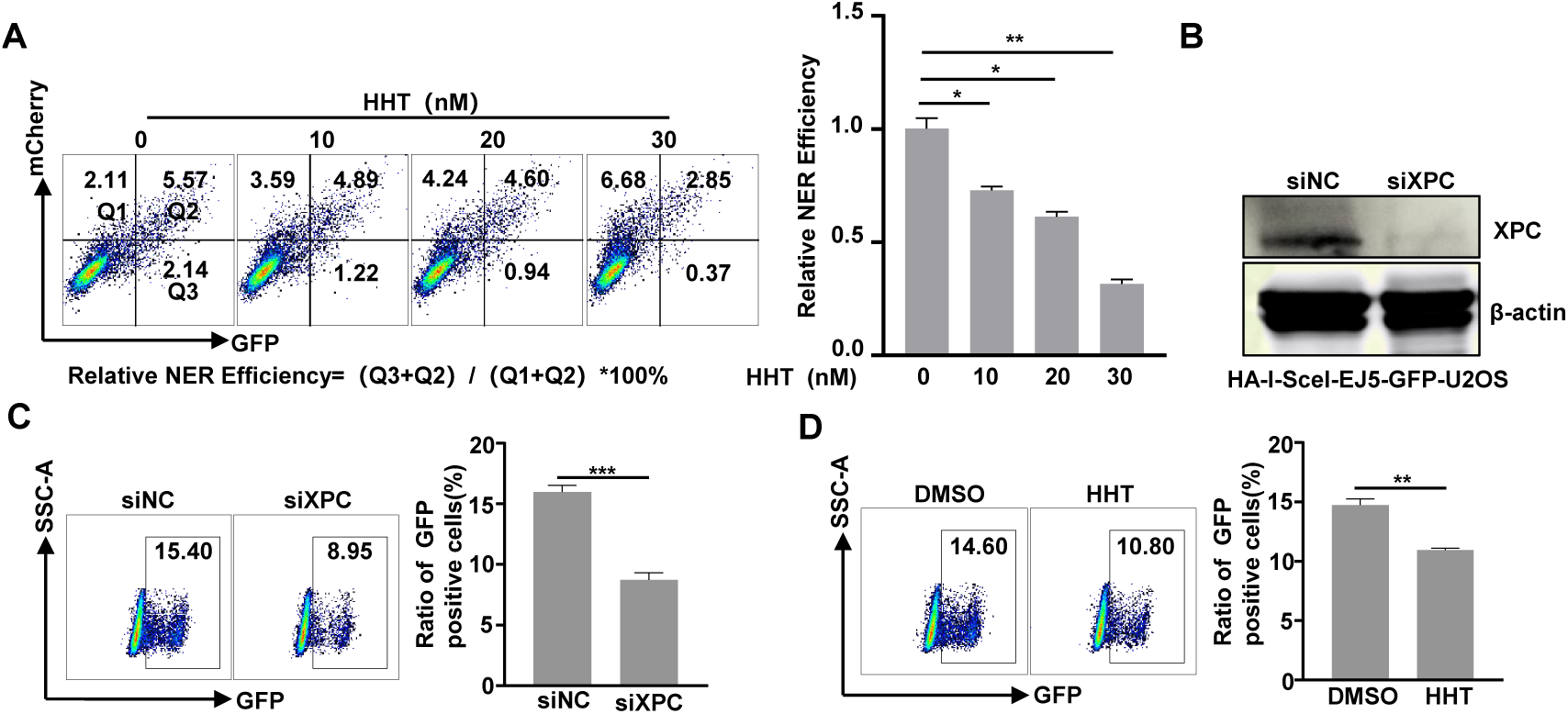
HHT impairs both NER and NHEJ efficiency. (A) Flow cytometry-based NER reporter assay in U2OS cells transfected with UV-damaged pEGFP-C3 plasmid and treated with increasing concentrations of HHT. Representative dot plots show GFP and mCherry signals used to calculate NER efficiency.Quantification of relative NER efficiency [(GFP/mCherry) normalized to control] across HHT treatment groups, demonstrating a dose-dependent reduction in NER activity upon HHT treatment. (B) Western blot analysis confirming XPC depletion in HA-I-SceI-EJ5-GFP-U2OS cells transfected with control (siNC) or XPC-targeting siRNA (siXPC). (C-D) Representative flow cytometry plots and quantification of GFP-positive cells showing that HHT treatment or XPC knockdown both significantly reduce NHEJ efficiency.

Although XPC is classically recognized as a core factor in NER, whether XPC participates in other DNA repair pathways remains unclear. Given that HHT markedly inhibited 53BP1 Tudor foci formation, a downstream marker of DNA double-strand break (DSB) repair signaling, we investigated whether XPC regulates DSB repair efficiency. To directly evaluate the role of XPC in NHEJ, we employed the doxycycline-inducible EJ5-GFP reporter system in U2OS cells, in which I-SceI-induced DNA double-strand breaks are repaired through NHEJ and quantified by restoration of GFP expression^32^ .Efficient depletion of XPC was confirmed by western blotting in HA-I-SceI-EJ5-GFP-U2OS cells (Fig.5B). Knockdown of XPC significantly reduced the percentage of GFP-positive cells following I-SceI induction, indicating impaired NHEJ repair efficiency(Fig. 5C) ,suggesting that XPC regulated the NHEJ, which is consistent with previous report, although the underlying mechnism need further investigation^33^. Consistently, treatment with HHT (30 nM) significantly reduced GFP-positive cells compared with DMSO-treated controls(Fig. 5D), further suggesting that XPC is required for efficient NHEJ-mediated DSB repair.

### HHT potently sensitizes tumor cells to platinum-based chemotherapy *in vitro* and ***in vivo***

One of the key therapeutic applications of DDR inhibitors is to sensitize tumor cells to chemotherapy. Platinum-based drugs are widely used as chemotherapeutic agents for multiple cancers, including triple-negative breast cancer (TNBC). These agents exert antitumor effects primarily by forming DNA intrastrand and interstrand crosslinks, and the resulting lesions are predominantly repaired via the nucleotide excision repair (NER) pathway. We therefore investigated whether HHT sensitizes triple-negative breast cancer MDA-MB-231 cells to platinum-based drugs.

As shown in Fig. S6,The IC₅₀ values of HHT against MDA-MB-231 was 44.43 nM . MDA-MB-231 cells were treated with a fixed low concentration of HHT (5 nM) in combination with increasing concentrations of cisplatin and carboplatin. Compared with platinum monotherapy, co-treatment with HHT markedly enhanced cellular sensitivity to cisplatin and carboplatin, with fold-increases of 4.8 and 4.3 respectively (Fig. 6A), indicating that HHT potently sensitizes tumor cells to platinum-based chemotherapy.

**Fig. 6.**
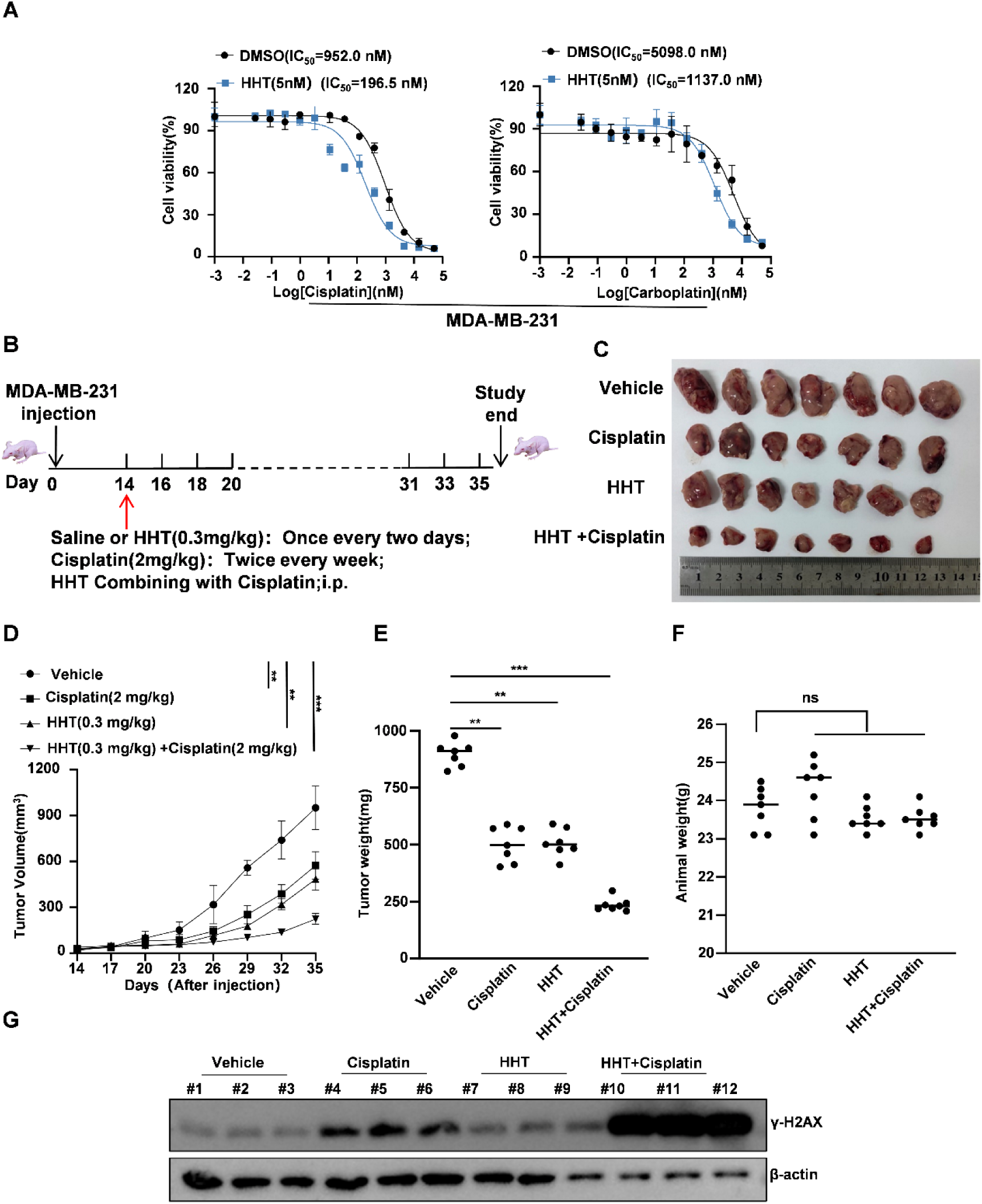
HHT enhances the antitumor efficacy of platinum-based chemotherapy in vitro and in vivo. (A) Dose–response curves showing cell viability of MDA-MB-231, SUM149PT, and MDA-MB-436 cells treated with Cisplatin or Carboplatin in the presence or absence of low-dose HHT (5 nM). Combination treatment shifts the dose–response curves leftward, indicating increased sensitivity to platinum-based drugs. (B) Schematic illustration of the drug administration schedule and experimental procedure in the mouse xenograft model. (C) Tumor growth curves of mice treated with vehicle, cisplatin, HHT, or the combination of HHT and cisplatin during the treatment period. (D) Representative images of tumors harvested at the experimental endpoint. (E) Tumor weights measured at the experimental endpoint. (F) Body weights of mice in each treatment group during the experiment. (G) Western blot analysis of γ-H2AX expression in tumor tissues from different treatment groups.

To determine whether HHT enhances the antitumor activity of cisplatin in vivo, we established an MDA-MB-231 cell-derived subcutaneous xenograft model, with the dosing regimen and treatment timeline shown in Fig. 6B. Both HHT and cisplatin as monotherapies suppressed tumor growth, tumor size and tumor weight, whereas their combination produced a significantly greater inhibitory effect than either agent alone(Fig. 6C–E). Importantly, no significant differences in mouse body weight were observed among all treatment groups (Fig. 6F).To further assess whether the enhanced antitumor effect correlated with increased DNA damage, tumor tissues were randomly collected from multiple mice per group and analyzed for γ-H2AX expression by western blotting. The combination group exhibited substantially higher γ-H2AX levels than the vehicle control or monotherapy groups (Fig. 6G). Collectively, these in vivo findings demonstrate that HHT enhances the antitumor efficacy of cisplatin, suppresses tumor proliferation, and promotes DNA damage accumulation in tumor tissues.

## Discussion

In this study, we demonstrate that HHT is a first-in-class XPC inhibitor that disrupts XPC–damaged DNA binding, impairs NER and NHEJ, and sensitizes tumor cells to platinum-based chemotherapy, offering a promising strategy to overcome platinum resistance.

HHT has been used clinically in China for more than 40 years and has shown favorable therapeutic efficacy against several hematological malignancies, including chronic myelogenous leukemia (CML) and acute myelogenous leukemia (AML)^34^. The canonical antileukemic mechanism of HHT involves binding to the A-site of the ribosome, thereby inhibiting protein translation^35^. Recent studies have further revealed the mechanistic diversity of HHT in AML by binding to NF-κB repressing factor (NKRF) ^27^and by modulating the phase separation of EWSR1^28^. Despite the well-documented clinical efficacy of HHT in the treatment of leukemia^36^, its application in solid tumors remains severely constrained by several critical bottlenecks, including undefined molecular targets, obscure mechanisms of action, and a lack of reliable predictive biomarkers^37^. Although a substantial body of evidence has demonstrated that HHT exerts growth-inhibitory effects on a broad spectrum of solid tumor cell lines— including those derived from breast, lung, colorectal, and liver cancers^38–41^—most current studies remain confined to the level of cell signaling pathways. Consequently, the direct targets and the precise mechanisms underlying the antitumor activity of HHT in solid tumors have yet to be fully elucidated. Our work identified XPC as a novel target of HHT in solid tumors. Moreover, we demonstrated that HHT disrupts the XPC– damaged DNA interaction and impairs DNA damage repair, including both NER and NHEJ(Fig.5). Therefore, we uncovered a previously unrecognized target and mechanism of HHT in solid tumor. More importantly, we found that HHT sensitizes tumor cells to platinum-based chemotherapy in vitro and in vivo, suggesting that HHT may be exploited to enhance platinum-based therapy in solid tumors.

Although XPC has been traditionally characterized as the primary damage recognition factor responsible for initiating global genome nucleotide excision repair (GG-NER)^42,43^, increasing evidence suggests that XPC possesses functions beyond canonical lesion repair^44^. Previous studies have indicated that depletion of XPC affects cellular responses to DNA damage beyond its established role in NER^33,45^, suggesting that XPC may function as a broader DDR regulator rather than merely an NER initiation factor. For example, XPC deficiency prevents recruitment of RNF8 to UV-induced DNA lesions^29^, and XPC knockdown cells exhibited an altered spectrum of NHEJ products with decreased levels of intramolecular joined products^33^.Our results demonstrate that bleomycin-induced 53BP1 Tudor foci formation is abolished in XPC-deficient cells, and that reintroduction of XPC restores Tudor foci formation upon bleomycin exposure; this restored foci formation is again inhibited by HHT treatment(Fig.3), further suggesting that XPC is an NHEJ regulator. Consistent with this, HHT also inhibits NHEJ through targeting XPC. In contrast, triptolide, an XPB inhibitor and well-established transcription-coupled NER inhibitor, does not inhibit NHEJ, suggesting that XPC regulates NHEJ independently of its NER function Fig. S7.

## METHOD

### Cell culture

HEK-293T,U2OS and MDA-MB-231 were purchased from ATCC and cultured in DMEM supplemented with 10% fetal bovine serum (FBS) at 37℃ in 5% CO_2_.

### DNA transfection, virus packaging and lentiviral infection

Cells were transfected using Polyethylenimine HCl MAX (Polysciences). Lentiviral supernatant, collected at 72 hours after co-transfection of vectors and packaging plasmids (pMD2G and psPAX2), was used for infection in the presence of 8 μg/mL polybrene. Stable transformants were then selected with 2 μg/mL puromycin over 4 days, and stability was confirmed via immunoblotting.

### High-throughput compound screening workflow

A compound library containing approximately 2592 compounds was screened using the EGFP-Tudor-U2OS reporter system. Screening was performed using 384-well plates, with each plate containing internal controls, including: Bleomycin-induced DNA damage control; ATM inhibitor plus Bleomycin treatment control. Compounds(10 μM) were added using an automated liquid-handling system. After 1 h compound incubation, Bleomycin(2 μM) was added to induce DSBs. Following 2 h DNA damage induction, plates were subjected to high-content imaging. To minimize variation caused by Bleomycin exposure time during sequential plate processing, compound addition and Bleomycin treatment were staggered, and the induction time for each plate was carefully recorded to maintain consistent DNA damage induction duration.

### Immunofluorescent staining

For immunofluorescence staining, cells were first fixed and permeabilized using 4% paraformaldehyde and 0.5% Triton X-100, respectively. Subsequent to blocking with 5% BSA in PBS, overnight incubation at 4°C was carried out with primary antibodies (γH2AX and 53BP1, each diluted 1:1000). Cells were then thoroughly washed with cold PBS (three times, 15 minutes each) prior to a 1hour incubation at room temperature with species-matched secondary antibodies conjugated to either Alexa Fluor 594.After repeating the washing procedure, nuclei were stained with DAPI. Imaging was performed on a fluorescence microscope following the application of an anti-fade mounting medium. Data analysis involved a t-test comparing at least 100 cells per condition, with results expressed as mean ± standard deviation (NS indicates no significant difference

### Cell viability assay

In 96-well plates, cells (300-500/well; triplicates per group) were first incubated for 24 h, then treated with HHT(out of toxic dose) and exposed to serial dilutions of Cisplatin , carboplatin or oxaliplatin for 7 days. The medium was then added with CellTiter-Glo Luminescent Cell Viability Assay; after 10 min of incubation, absorbance was measured at 450 nm to assess viability.

### Protein extraction and western blotting

U2OS were transfected with FLAG-XPC(WT), FLAG-XPC-TST3A, FLAG-XPC-T117A, FLAG-XPC-S122A and FLAG-XPC-T795A. Equal cell numbers were scraped, pelleted, and lysed on ice for 25 min in NETN buffer (20 mM Tris-HCl pH 8.0, 100 mM NaCl, 1 mM EDTA, 0.5% Nonidet P-40) with protease inhibitors. After clearing by centrifugation (12,000 rpm, 10 min),80 μL of lysate was mixed with 20 μL of 5× SDS loading buffer and boiled (99°C, 10 min) as input. The remaining lysate was incubated with NCT/HHT-Beads at 4°C for 2 h; beads were then washed with NETN buffer, boiled in 1× SDS loading buffer, and the eluate was separated by SDS-PAGE. Proteins were transferred to Immobilon®-FL PVDF membranes (Millipore), blocked with 5% milk/TBST, and immunoblotted with the indicated antibodies.

### Identification of HHT-interacting proteins by IP/MS

U2OS cell lysates were incubated with HHT-conjugated beads or control beads overnight at 4 °C. After extensive washing, specifically bound proteins were eluted and subjected to LC–MS/MS analysis. Raw data were analyzed using MaxQuant against the human UniProt database. Candidate proteins were identified based on enrichment in HHT probe pull-down compared with control beads.

### CRISPR/Cas9-mediated gene knockout

Gene knockout cell lines were generated using the CRISPR/Cas9 genome-editing system. Single-guide RNAs (sgRNAs) targeting the coding region of the target gene were designed using the CRISPR design tools (CRISPOR) and selected based on high on-target activity and minimal predicted off-target effects.The sgRNA sequences were cloned into a Cas9-expressing lentiviral vector (such as lentiCRISPR v2) containing both the Cas9 nuclease and sgRNA expression cassette. Lentiviral particles were produced by co-transfection of the sgRNA-Cas9 plasmid together with packaging plasmids into HEK293T cells.Target cells were infected with lentivirus expressing target gene-specific sgRNA or a non-targeting control sgRNA. Following infection, cells were selected with appropriate antibiotics to enrich for transduced populations. Single-cell clones were subsequently obtained by limiting dilution or fluorescence-activated cell sorting (FACS).

### Expression and purification of recombinant proteins

The XPC was reconstituted and purified using the following procedure. A 0.8 L culture of Sf9 insect cells expressing His-tagged XPC/XPC-T795A was pelleted using lysis buffer (50 mM Tris-HCl, pH 7.5, 300 mM KCl, 20 mM imidazole, 5% glycerol, 10 mM β-mercaptoethanol, 0.01% Triton X-100) supplemented with protease inhibitors (0.25 mM PMSF, 10 µg/ml pepstatin A, and one EDTA-free Complete protease inhibitor cocktail tablet from Roche). The mixture was lysed via sonication. The resulting lysate was clarified by centrifugation at 35,000 × g for 30 min at 4°C and then applied to a 1 ml Ni NTA Beads (Smart-lifesciences). Following a wash with lysis buffer containing 1 M KCl, the complex was eluted using lysis buffer with 300 mM imidazole. The eluted XPC-containing fractions were concentrated and subjected to size-exclusion chromatography on a Superose 6 column (10/300, GE Healthcare) pre-equilibrated with storage buffer (25 mM Tris-HCl, pH 7.4, 300 mM KCl, 5 mM DTT, 10% glycerol, 0.007% Triton X-100). The peak fractions corresponding to the XPC were pooled, concentrated, and stored at –80°C.

### EMSA assay

The Bot56 DNA strand was 5’-end labeled with biotin and subsequently annealed with its complementary Top strand (Top56_2AP) to generate lesion-containing duplexes. For binding reactions, 5 to 10 nM of the labeled DNA duplex was incubated with an equimolar amount of XPC mixed with recombinant SBP-Rad23B and SBP-Centrin2, in a 20 μl volume of reaction buffer (25 mM Tris-HCl, pH 8.0, 100 mM KCl, 8% glycerol, 0.007% Triton X-100, 5 mM DTT, 7 mM MgCl₂) at 30°C for 30 minutes. The reaction mixtures were then resolved on a 4% native polyacrylamide gel in 1X TBE buffer at 4°C. Gels were imaged using a Typhoon FLA 9500 Phosphor Imager.

### Flow Cytometry

Flow cytometric analysis was conducted to evaluate the efficiency of NER. Treated and control cells were harvested, washed with PBS, All stained samples were analyzed with a BD LSRFortessa cell analyzer (BD Bioscience, USA). A minimum of 10,000 gated events per sample were collected. Data analysis, including compensation and gating based on FSC/SSC and single-stain controls, was performed using FlowJo 10.8.1.

### Cellular Thermal Shift Assay (CETSA)

Target engagement of HHT was assessed by CETSA. U2OS, treated with compound or vehicle control, were harvested, washed in PBS with protease inhibitors, and aliquoted. Aliquots were heated at a defined temperature gradient (37–61°C) for 3 min in a thermal cycler, followed by room temperature incubation for 3 min. Cells were lysed by three freeze-thaw cycles (liquid nitrogen/25°C water bath). Lysates were cleared by centrifugation (20,000 × g, 20 min, 4°C). Soluble protein was quantified (BCA assay), separated by SDS-PAGE, and immunoblotted for the protein of interest. Band intensities were quantified (ImageJ), and the percentage of soluble protein relative to the 37°C control was plotted against temperature to generate melting curves.

### Molecular docking

The molecular docking procedure was executed using the AutoDock Vina 2.1.6 software. From the ensemble of generated docking poses, the representative conformation was chosen according to two primary criteria: possessing the lowest binding energy score and being the most frequently recurring pose among high-ranking solutions. This selected pose was then imported into PyMOL 3.1 and Discovery Studio 2019 for visualization and further interaction analysis.

### Cell-derived xenograft models and drug treatment

All animal experiments were approved by the Institutional Animal Care and Use Committee of Chongqing Medical University. Five-week-old female BALB/c nude mice were used to establish xenograft models. Briefly, 3 × 10⁶ MDA-MB-231 cells suspended in Matrigel/PBS mixture were subcutaneously injected into the left anterior axillary region of mice. When tumors reached approximately 80–100 mm³, mice were randomly divided into four groups: vehicle control, HHT, cisplatin, and HHT plus cisplatin groups. HHT was administered intraperitoneally at 0.3 mg/kg every two days, whereas cisplatin was administered intraperitoneally at 2 mg/kg twice weekly. Treatment continued for 35 days. Tumor volume was measured every 2–3 days using calipers and calculated as (length × width²)/2. Mice were euthanized when tumor volume reached the ethical endpoint (≤1200 mm³). Tumors and major organs were collected for further analysis.

### Statistical analysis

Data analysis was performed with GraphPad Prism 10. Results are shown as mean ± SEM(n ≥ 3 independent experiments). Significance between two groups was determined by two-tailed t-tests; for multiple groups, one-way or two-way ANOVA was used as appropriate (see legends). A *p*-value < 0.05 was considered statistically significant.

## Authors contributions

Jiaxue Wu and Yongjun Dang conceived and designed the study. The order of the co–first authors’ names was determined on the basis of their contributions to this study (Chuanjie Wang and Ying Zhang performed a majority of the experiments, and Xiaoqin Zhou completed the animal experiments and analyzed the related data). Jia He provided HHT-Beads, Longjiang Liu prepared EGFP-Tudor-U2OS. Luyi Wang and Yanfang Zhang performed the immunohistochemistry experiments, Qin Gong predicted the binding site of HHT and XPC, Jiayu Zhang performed Western blot analysis. Chuanjie Wang wrote the manuscript. Jiaxue Wu and Yongjun Dang edited the manuscript. All authors read and approved the final manuscript.

## Data availability

The data supporting the findings of this study are available from the corresponding authors upon reasonable request.

## Dual publication

The authors confirm that the manuscript has not been submitted to or published in any other journal simultaneously.

## Acknowledgements

The animal experiment in this paper was partly supported by the Animal Experimentation Center of Chongqing Medical University

